# Sirtuin3 ensures the metabolic plasticity of neurotransmission during glucose deprivation

**DOI:** 10.1101/2023.03.08.531724

**Authors:** Anupama Tiwari, Arsalan Hashemiaghdam, Marissa A. Laramie, Dario Maschi, Tristaan Haddad, Marion I. Stunault, Carmen Bergom, Ali Javaheri, Vitaly Klyachko, Ghazaleh Ashrafi

## Abstract

Neurotransmission is an energetically expensive process that underlies cognition. During intense electrical activity or dietary restrictions, glucose levels in the brain plummet, forcing neurons to utilize alternative fuels. However, the molecular mechanisms of neuronal metabolic plasticity remain poorly understood. Here, we demonstrate that glucose-deprived neurons activate the CREB and PGC1α transcriptional program that induces the expression of the mitochondrial deacetylase Sirtuin 3 (Sirt3) both *in vitro* and *in vivo*. We show that Sirt3 localizes to axonal mitochondria and stimulates mitochondrial oxidative capacity in hippocampal nerve terminals. Sirt3 plays an essential role in sustaining synaptic transmission in the absence of glucose by powering the retrieval of synaptic vesicles after release. These results demonstrate that the transcriptional induction of Sirt3 ensures the metabolic plasticity of synaptic transmission.

**Highlights:**

- Glucose deprivation drives transcriptional reprogramming of neuronal metabolism via CREB and PGC1α.
- Glucose or food deprivation trigger the neuronal expression of mitochondrial deacetylase sirtuin 3 (Sirt3) both *in vitro* and *in vivo*.
- Sirt3 stimulates oxidative ATP synthesis in nerve terminals.
- Sirt3 sustains the synaptic vesicle cycle in the absence of glucose.

## Introduction

The brain requires a constant and ready source of energy to function properly. Indeed, cognitive function is particularly susceptible to metabolic perturbations, such as, ischemic stroke or hypoglycemia. Glucose is considered the primary fuel for the brain, yet glucose concentration in the interstitial fluid (ISF) surrounding neurons is considerably lower (0.5-1mM in rodents) than in the blood^1–3^. Furthermore, the glucose content of ISF is locally depleted in brain regions during bouts of neuronal activity^4, 5^. Availability of glucose in the brain is further exacerbated by the relative scarcity of glucose storage as glycogen^6^, forcing neurons to rely on alternative energy sources, such as, lactate, ketone bodies, and amino acids, during hypoglycemia^7–9^. Lactate, and its derivative pyruvate, are major neuronal fuels that are supplied systemically through the circulation or by local astrocytic supply. Indeed, astrocytes have been shown to provide neurons with lactate produced from glutamate and other amino acids in a process known as the astrocyte-to-neuron lactate shuttle^10, 11^. Lactate and pyruvate are oxidative fuels that bypass glycolysis and are directly broken down via mitochondrial oxidative phosphorylation (OXPHOS). Therefore, in the absence glucose, neurons need to increase their mitochondrial capacity for OXPHOS to compensate for lack of glycolytic ATP production. However, the mechanisms of neuronal metabolic adaptation to low glucose and their physiological impact on synaptic function remain poorly understood.

In most cell types, metabolic adaptation to energetic stress, such as fuel shortage, is initiated transcriptionally through coordinated control of gene expression. It is well known that neuronal electrical activity or muscle contraction activate the transcription factor family of cAMP-response element-binding proteins (CREBs)^12–14^. One of the target genes of CREB transcription program is the transcriptional coactivator PGC1α (peroxisome proliferator-activated receptor gamma coactivator-1 alpha), which plays a key role in mitochondrial biogenesis and OXPHOS. In skeletal muscle, caloric restriction and exercise induce the expression of PGC1α^15^ and its transcriptional target Sirtuin 3 (Sirt3), a mitochondrial lysine deacetylase enzyme^16^. Sirt3 resides in the mitochondrial matrix that regulates mitochondrial metabolism^17^ and resistance to oxidative stress^18^ in diverse cell types. Sirt3 is one of the most abundant sirtuins in the brain^19^,but its function in the metabolic support of synaptic transmission remains unexplored.

In neurons, nerve terminals are loci of high energy demand due to their role in synaptic transmission. We and others have previously shown that during brief periods of glucose deprivation, mitochondrial oxidation of pyruvate can sustain the synaptic vesicle (SV) cycle^20, 21^, a crucial process in synaptic transmission. In the present study, we examined the metabolic adaptations of neurons to prolonged glucose deprivation and revealed a critical role for the mitochondrial deacetylase Sirt3 in ensuring the metabolic plasticity of synaptic transmission.

## Results

### Neuronal glucose deprivation induces transcriptional reprogramming of mitochondrial metabolism

We previously demonstrated that neurons utilize the oxidative fuels lactate and pyruvate to support synaptic transmission during acute glucose deprivation^20^. Other studies have also shown that neuronal activity continues to persist during prolonged glucose shortage in fasting or restrictive diets^22^ implicating yet unknown mechanisms of metabolic plasticity in neurons. We examined the transcriptional rewiring of neuronal metabolism during glucose deprivation by performing RNA sequencing on primary cultures of cortical neurons maintained in glucose-rich or glucose-depleted (and serum-free) media (**Figure 1A**). Pathway analysis of differentially expressed genes revealed the induction of genes involved in the unfolded protein response mediated by PERK, ATF4, and ATF6 transcription factors (**Figure 1B**). Furthermore, plasma membrane amino acid transporters were upregulated under glucose deprivation, suggesting the reprogramming of metabolic pathways to utilize amino acids as an alternative energy source^23^.

**Fig 1.**
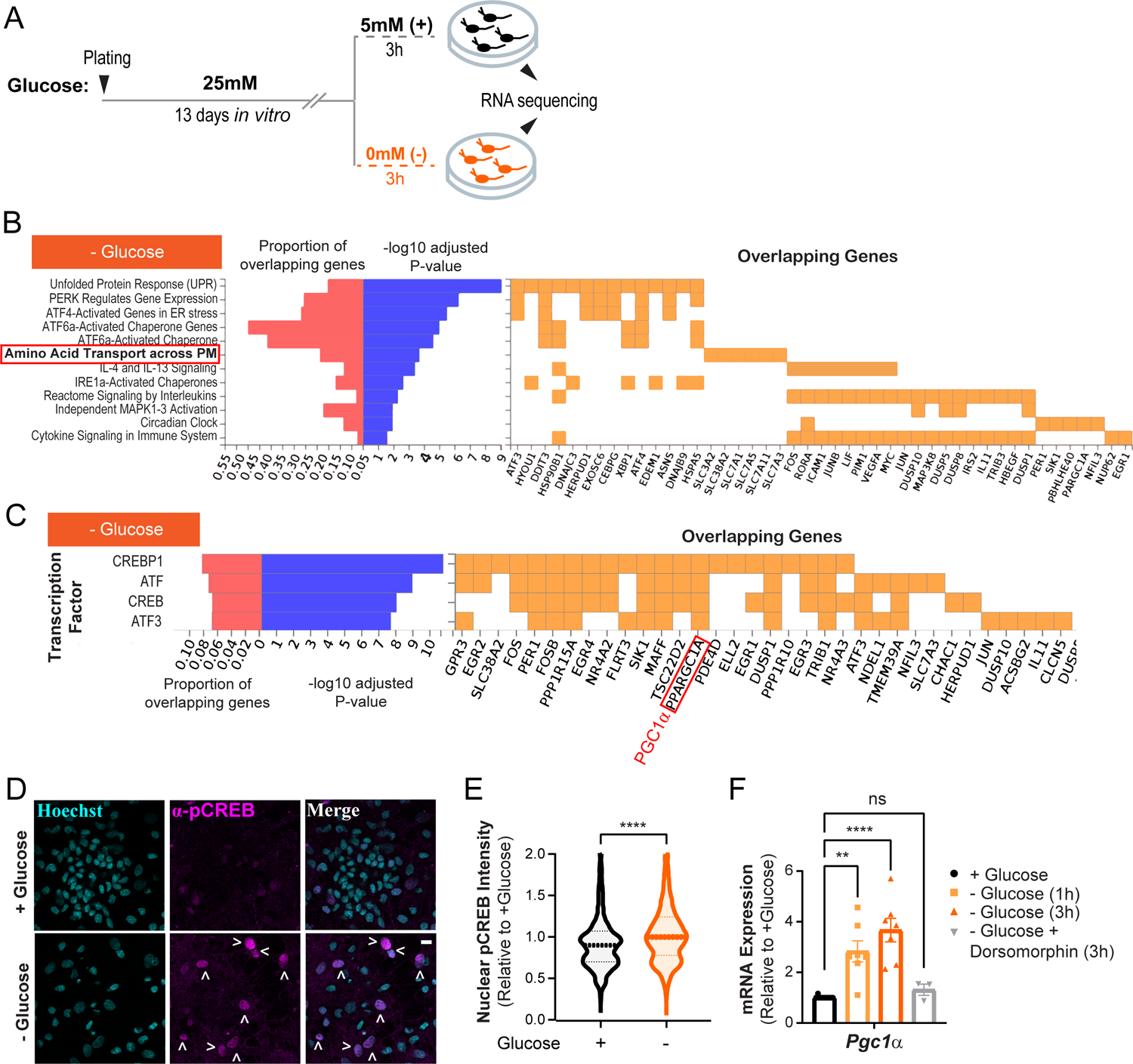
Transcriptional reprogramming of neuronal metabolism during glucose deprivation. **A)** A paradigm for glucose deprivation of primary cortical neuronal cultures. **B)** Pathway analysis of differential gene expression in neurons after glucose deprivation (a selection of top pathways and genes are shown). **C)** Gene targets of the CREB transcription factors are enriched in the glucose-deprived dataset. The target gene Pgc1α, denoted as PPARGC1A, is boxed in red. **D)** Immunostaining of cortical neuronal cultures (treated as in A) with an anti-phospho-CREB antibody and Hoechst nuclear stain. Arrowheads denote pCREB-positive nuclei. **E)** Fluorescence intensity of nuclear phospho-CREB staining relative to the +glucose (control) condition. Normalized intensity ± SEM: + glucose (3h), 1.00 ± 0.01, - glucose (3h), 1.13 ± 0.01. n= 4046 - 6925 nuclei. **F)** Relative mRNA expression of *Pgc1α* in neuronal cultures treated as in panel A, with or without the AMPK inhibitor, dorsomorphin. Values are normalized to *β-actin* mRNA and expressed relative to the +glucose condition. n=3-10 cortical samples. Average normalized mRNA level ± SEM: - glucose (1h), 2.83 ± 0.42; - glucose (3h), 3.67 ± 0.46; - glucose + dorsomorphin (3h), 1.32 ± 0.22. **p < 0.01 and ****p < 0.0001; One-way ANOVA (E), Mann-Whitney U test (F). Violin plot represents median 25^th^ and 75^th^ percentiles (dotted lines) with outliers removed. The bar graph is plotted as mean ± SEM. Table. S1. (Related to Fig.1) List of differentially expressed genes in glucose-deprived neurons.

Glucose deprivation also stimulated the expression of genes in the CREB transcription program (**Figure 1C**), which control metabolic homeostasis^14^. We confirmed the activation of CREB pathway by the immunostaining of primary cortical cultures with an anti-phospho-CREB antibody, which revealed elevated nuclear phospho-CREB staining during glucose deprivation (**Figures 1D and 1E**). Amino acids are oxidized inside the mitochondria and they enter the tricarboxylic acid cycle as pyruvate, oxaloacetate, or α-ketoglutarate. Indeed, we found that PGC1α (also known as PPARGC1A), the master regulator of mitochondrial energy metabolism^24^, was among the CREB target genes that were induced during glucose deprivation (**Figure 1C**, boxed). The induction of *Pgc1α* mRNA during glucose deprivation was also validated with quantitative real-time PCR (qPCR) (**Figure 1F**).

Energetic stress in most cell types activates the cellular energy sensor, AMP kinase, by increasing the AMP (or ADP) to ATP ratio^25–27^. In turn, AMP kinase initiates pathways of metabolic adaptation, including the CREB transcriptional program^28^. Consistent with this upstream regulatory role, we found that inhibition of AMP kinase with dorsomorphin^29^ blocked the induction of *Pgc1α* expression in glucose-deprived neurons (**Figure 1F**), indicating that this transcriptional response is driven by energetic stress. We conclude that neuronal glucose deprivation triggers the CREB transcriptional program resulting in activation of PGC1α, a potent regulator of mitochondrial oxidative metabolism^30^.

### Glucose deprivation stimulates the expression of Sirtuin 3 and deacetylation of mitochondrial proteins

Post-translational protein modifications play a critical role in rapid and reversible modulation of cellular homeostasis. In calorically restricted skeletal muscle, the transcription coactivator PGC1α stimulates the expression of the mitochondrial deacetylase enzyme Sirt3 by binding to its promoter region^31^. Sirt3 drives metabolic reprogramming in muscle by deacetylating mitochondrial proteins involved in OXPHOS^17, 32^. The induction of PGC1α (**Figure 1F**) in glucose-deprived neurons prompted us to investigate whether Sirt3 expression was similarly upregulated. Indeed, *sirt3* mRNA level was elevated in glucose-deprived cortical neurons, and this induction was blocked by the pharmacological inhibition of AMP kinase (**Figure 2A**). Furthermore, we found that adenoviral overexpression of PGC1α^33^ in cortical neurons was sufficient to induce *sirt3* gene expression even in the presence of glucose, implicating PGC1α as a transcriptional regulator of neuronal Sirt3 (**Figure 2B**). In agreement with its mRNA induction, the expression of Sirt3 protein was similarly elevated in glucose-deprived neurons (**Figures 2C and 2D**).

**Fig 2.**
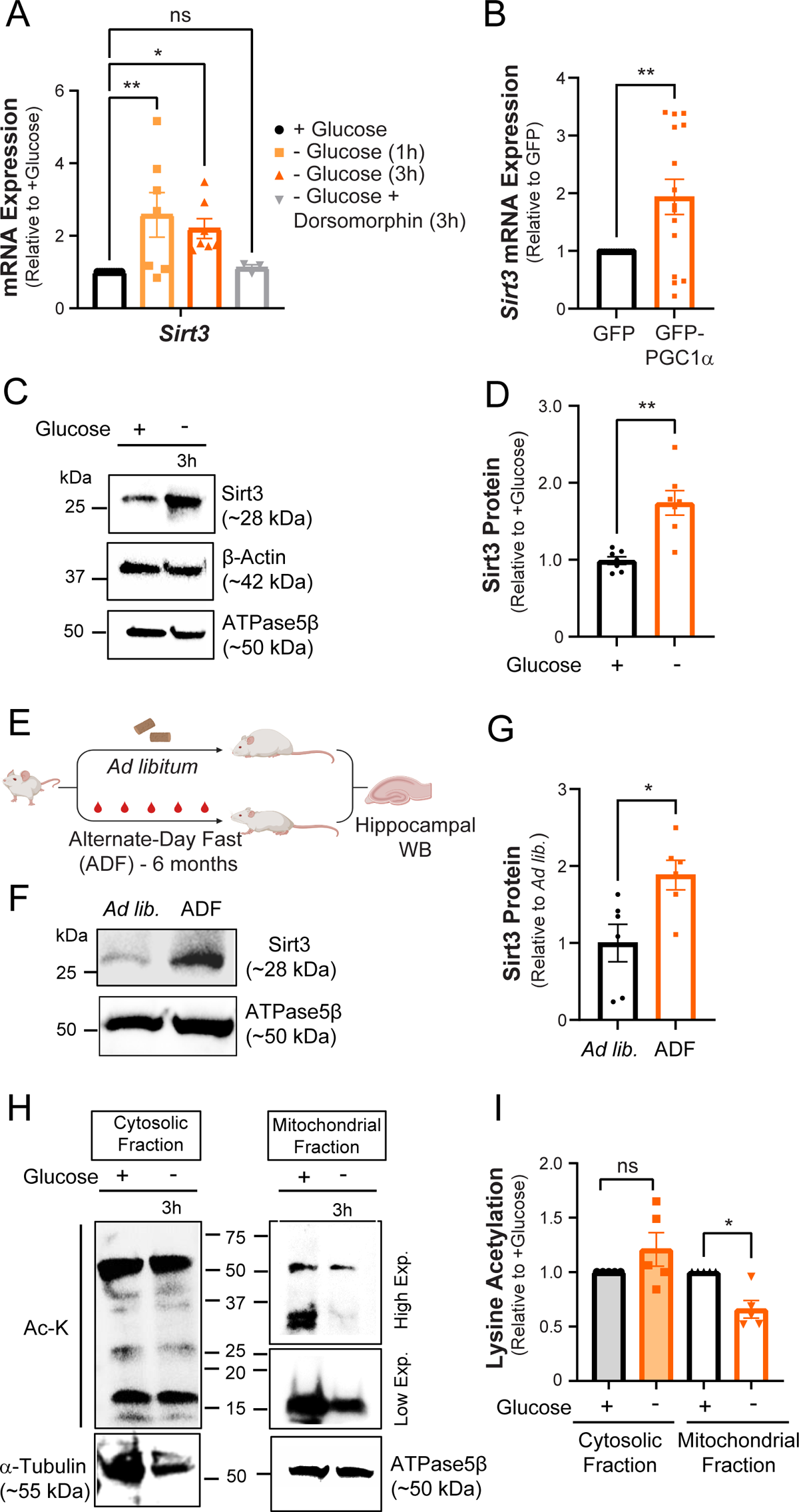
Glucose deprivation stimulates neuronal Sirt3 expression and deacetylation of mitochondrial proteins. **A)** Relative *Sirt3* mRNA expression in control and glucose-deprived neurons. Values are normalized to *β-actin* mRNA and expressed relative to control (+glucose). n = 3-10 cortical samples. Average normalized mRNA level ± SEM: - glucose (1h), 2.58 ± 0.62; - glucose (3h), 2.20 ± 0.27; - glucose + dorsomorphin (3h), 1.12± 0.08. **B)** Relative *Sirt3* mRNA expression in cultures transduced with adenoviral particles encoding GFP (control) and GFP-PGC1α, normalized to *18s* rRNA and expressed relative to control. n=14 cortical samples. Average normalized mRNA level ± SEM: GFP-PGC1α, 1.93 ± 0.30. **C)** Immunoblotting of Sirt3 protein expression in cortical neurons immunoblotted with antibodies against Sirt3 and the cytosolic and mitochondrial controls, β-actin, and ATPase5β, respectively. **D)** Sirt3 band intensity normalized to the ATPase5β and expressed relative to control. n = 7 cortical samples. Average normalized Sirt3 band intensity ± SEM: +glucose, 0.99 ± 0.05; - glucose, 1.74 ± 0.16. **E)** A paradigm for analysis of Sirt3 expression in hippocampi of mice fed *ad libitum* (*ad lib*) or alternate-day fasted for 6 months (ADF). **F)** Immunoblotting of Sirt3 protein in mouse hippocampal lysates. **G)** Sirt3 band intensity normalized to the ATPase5β band and expressed relative to the *ad lib* mice. n = 6 animals. Average normalized Sirt3 band intensity ± SEM: *ad lib*, 1 ± 0.24; ADF (6 months), 1.88 ± 0.19. **H)** Mitochondrial and cytosolic fractions isolated from cortical neuronal cultures maintained for 3 hours with (+) or without (-) of glucose were immunoblotted for acetylated lysines (Ac-K), α-tubulin (cytosolic marker) and ATPase5β. **I)** Intensity of lysine acetylation bands normalized to α-tubulin or ATPase5β plotted relative to control. n = 5 cortical samples. Average normalized Ac-K intensity: cytosolic fraction (-glucose), 1.21 ± 0.15; mitochondrial fraction (-glucose), 0.66 ± 0.08. *p < 0.05; one-way ANOVA (A), unpaired t-test (B), Mann-Whitney U-test (D, G, and I).

Given our *in vitro* findings, we investigated whether neuronal Sirt3 expression was similarly regulated by fuel availability *in vivo*. In agreement with a previous report^34^, we found that after 6 months of alternate-day fasting (ADF), the expression of Sirt3 in mouse hippocampi was elevated when compared to mice fed *ad libitum* (**Figures 2E-2G**). In contrast, overnight fasting did not alter hippocampal Sirt3 level (**Figures S1A and S1B**), suggesting that the regulation of Sirt3 expression is an adaptive response to fuel availability rather than a short-term stress response. Collectively, our data suggest that fuel deficiency stimulates neuronal Sirt3 expression both *in vitro* and *in vivo*.

Sirt3 abundance often correlates inversely with acetylation of mitochondrial proteins. This is exemplified by hyper-acetylation of mitochondrial proteins in *Sirt3^-/-^* tissues^16, 35, 36^, and deacetylation of mitochondrial proteins in exercising muscle with elevated Sirt3 expression^32^. The induction of Sirt3 in glucose-deprived neurons prompted us to quantitatively examine its effects on mitochondrial protein acetylation. To this end, mitochondrial fractions prepared from neuronal lysates were immunoblotted with an antibody against acetylated lysine residues, revealing a significant reduction in mitochondrial protein acetylation in cortical neurons deprived of glucose, while acetylation of cytosolic proteins remained unchanged (**Figures 2H and 2I**). Thus, we conclude that neuronal glucose deprivation stimulates Sirt3 expression downstream of PGC1α, thereby leading to deacetylation of mitochondrial proteins.

### Sirt3 is expressed in neuronal mitochondria and colocalizes with presynaptic terminals

Previous studies have shown that Sirt3 is expressed in mouse brain lysates from multiple regions, including, the cortex and hippocampus^19^. However, the subcellular distribution of Sirt3 in neuronal compartments remains unclear. Therefore, we performed immunocytochemistry on dissociated hippocampal neurons using antibodies against Sirt3 along with specific mitochondrial or neurite, or presynaptic markers. We found that Sirt3 localizes to neuronal mitochondria both in the soma and along neurites, as indicated by immunostaining for the mitochondrial marker TOMM20 (**Figure 3A**). Similar to the sparse distribution of mitochondria along neuronal axons^37^, Sirt3 is expressed along hippocampal neurites and colocalizes with presynaptic terminals, as indicated by co-staining with the neuritic marker Tuj-1 (β-tubulin III) (**Figure 3B**), and the synaptic marker vGLUT1 (**Figure 3C**). Our results thus confirm that Sirt3 localizes to neuronal axons, and is present at presynaptic terminals, suggesting a potential role for modulation of synaptic function.

**Figure 3.**
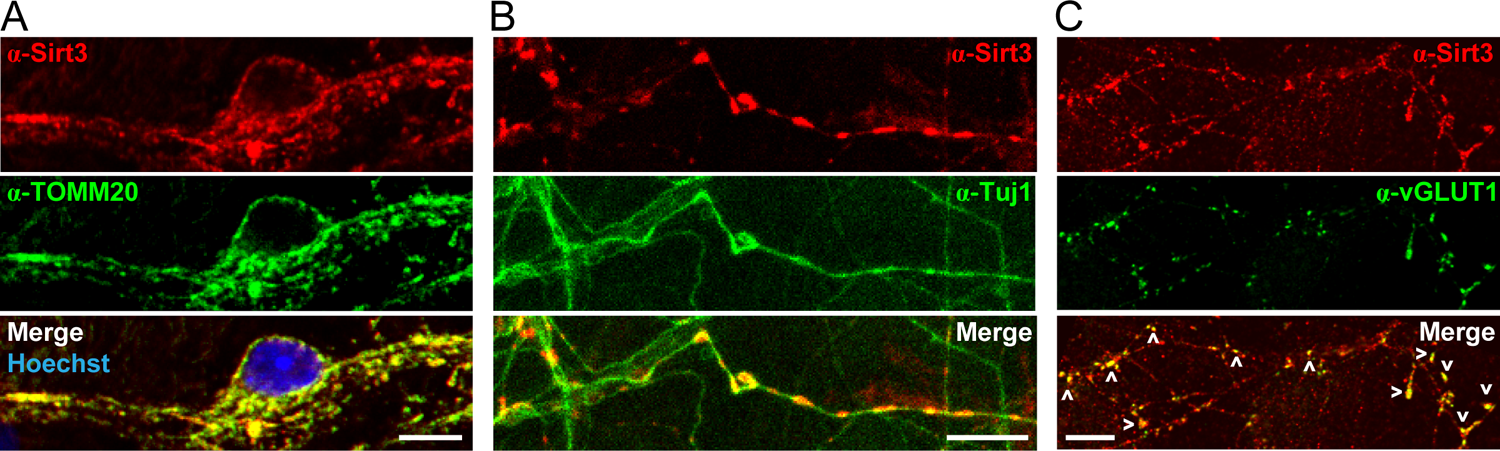
Sirt3 is expressed in neuronal mitochondria and colocalizes with presynaptic terminals. **(A-C)** Dissociated rat hippocampal neurons co-immunostained for Sirt3 and the mitochondrial marker TOMM20 **(A)**, the neurite marker Tuj1 **(B)**, or the synaptic marker VGLUT1 **(C)**, and the Hoechst nuclear stain **(A)**. Arrowheads indicate colocalization of presynaptic terminals with Sirt3 **(C)**. Scale bars, 10 μm.

### Sirt3 stimulates mitochondrial ATP synthesis and sustains synaptic transmission

Nerve terminals are loci of high energy demand and we have previously shown that synaptic function can be sustained by mitochondrial oxidation of pyruvate in the absence of glucose. Given that Sirt3 modulates mitochondrial metabolism in non-neuronal cells, and it is expressed in presynaptic mitochondria (**Figure 3C**), we examined the metabolic function of Sirt3 in terminals that were deprived of glucose. To this end, we monitored presynaptic ATP level in dissociated hippocampal neurons before, during, and after electrical stimulation using Syn-ATP, a genetically encoded ATP indicator targeted to nerve terminals^38, 39^ (**Figure 4A**). To mimic the metabolic shift to mitochondrial OXPHOS that occurs in response to glucose deprivation (**Figure 1**), neurons were supplied with a 1:1 mixture of lactate and pyruvate, replacing glucose. Compared to control neurons, presynaptic ATP level prior to electrical stimulation (pre-stimulation) was significantly lower in nerve terminals where Sirt3 was transiently knocked down with a short hairpin RNA (shRNA) construct, referred to as Sirt3 KD (**Figures 4B and 4C**). The efficiency of Sirt3 shRNA was confirmed by qPCR of *sirt3* mRNA level in cortical neuronal cultures (**Figure S2A**). Robust action potential (AP) firing (600 AP at 10 Hz) did not cause further ATP depletion, although post-stimulation ATP remained lower in Sirt3 KD terminals than in control ones (**Figures 4B and 4C**). These results demonstrate that Sir3 stimulates mitochondrial capacity for oxidative ATP production in nerve terminals.

**Fig 4.**
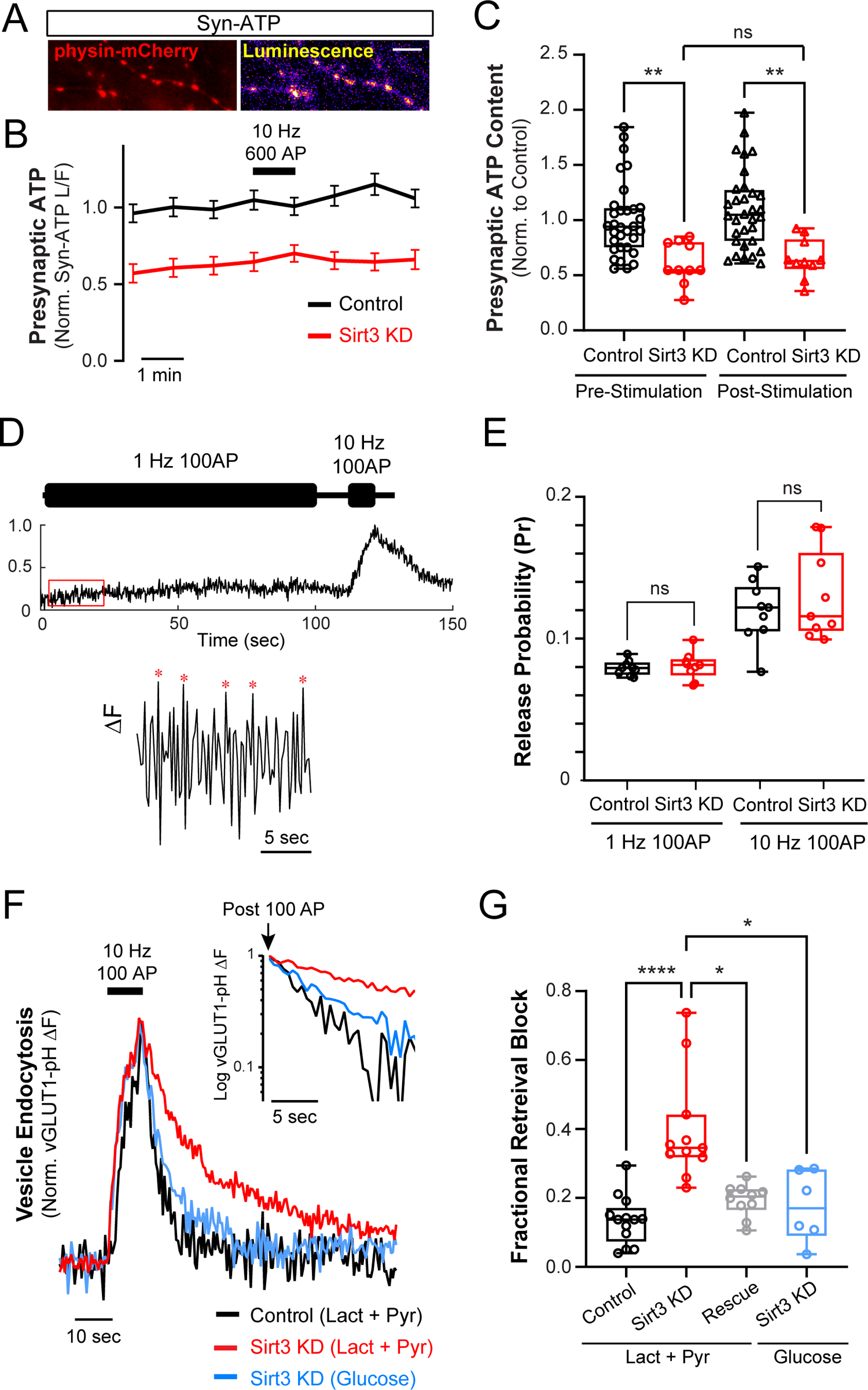
Sirt3 stimulates oxidative ATP production and sustains synaptic transmission in hippocampal nerve terminals. **A)** luminescence and physin-mCherry fluorescence images of hippocampal nerve terminals expressing Syn-ATP. **B)** Normalized presynaptic ATP traces in control and Sirt3 KD neurons supplied with lactate and pyruvate and stimulated with 600 AP at 10 Hz. **C)** Normalized ATP level before and after electrical stimulation. Average ATP level: control (pre-stimulation), 1.00 ± 0.06, Sirt3 KD (pre-stimulation), 0.61 ± 0.19, control (post-stimulation), 1.10 ± 0.35, Sirt3 KD (post-stimulation), 0.65 ± 0.18. n= 10-32 neurons. **D)** SV release events (indicated by asterisks) in terminals expressing vGLUT1-pH stimulated with 100 AP at 1 Hz, or at 10 Hz. **E)** SV release probability (P_r_) in control and Sirt3 KD terminals. Average P_r_ ± SEM: control (1Hz), 0.079 ± 0.005, Sirt3 KD (1 Hz), 0.081 ± 0.003, control (10 Hz), 0.112 ± 0.007, Sirt3 KD (10 Hz), 0.116 ± 0.011.n = 9 neuronal samples. **F)** Sample vGLUT1-pH traces of control, Sirt3 KD, and rescue terminals (expressing shRNA-resistant Sirt3) stimulated with 100 AP at 10 Hz in the presence of glucose or lactate/pyruvate. (Inset) semi-log plot of vGLUT1-pH traces following stimulation. **G)** Fractional retrieval block of vGLUT1-pH calculated as described in methods. n= 6-13 neurons. Average retrieval block ± SEM: control (lact+pyr), 0.14 ± 0.02, Sirt3 KD (lact+pyr), 0.40 ± 0.05, rescue (lact+pyr), 0.19 ± 0.01, Sirt3 KD (glucose), 0.18 ± 0.04. n.s., not statistically significant. Kruskal-Wallis test (C and G), Two-sample t-test (E). Black bars denote electrical stimulation. The box-whisker plots denote median (line), 25^th^-75^th^ percentile (box), and min-max (whiskers).

We previously demonstrated that mitochondrial Ca^2+^ uptake is essential for maintaining presynaptic ATP levels during electrical activity. To determine whether Sirt3 regulates mitochondrial Ca^2+^ dynamics, we monitored mitochondrial Ca^2+^ uptake in nerve terminals using a mitochondrially targeted GCaMP6 (mito^4xi^-GCaMP6f). Mitochondrial Ca^2+^ uptake in response to brief bursts of AP firing was not perturbed in Sirt3 KD terminals (**Figures S2C and S2D**), consistent with ATP level remaining stable post-stimulation (**Figure 4B**). Furthermore, we examined mitochondrial morphology with confocal microscopy revealing fragmentation of axonal mitochondria in Sirt3 KD neurons (**Figure S2B**), consistent with the role of Sirt3 in regulating mitochondrial fusion^40^. Altogether, Ca^2+^ and ATP measurements in nerve terminals demonstrate that Sirt3 stimulates neuronal mitochondrial capacity for oxidative ATP production without altering mitochondrial Ca^2+^ dynamics during electrical stimulation.

Several studies have shown that mitochondrial oxidation of lactate can replace glucose to power energy-sensitive steps in the SV cycle^20, 21^. Given that Sirt3 is essential for mitochondrial ATP synthesis in nerve terminals, we investigated the effects of Sirt3 depletion on the SV cycle in the absence of glucose (**Figure 4D-G**). We monitored AP-driven release, endocytosis, and re-acidification of SVs using vGLUT1-pHluorin (vGLUT1-pH) in nerve terminals supplied with lactate and pyruvate but not glucose. We first examined SV release in terminals that were electrically stimulated with trains of 100 APs at low (1 Hz) or high (10Hz) activity levels (**Figure 4D**). We determined that Sirt3 KD did not alter SV release probability at 1 Hz or 10 Hz stimulation as compared to control (**Figure 4E**). In contrast, SV endocytosis was significantly less efficient in Sirt3 KD terminals supplied with lactate and pyruvate, but this defect could be rescued by expression of an shRNA-resistant Sirt3 construct (**Figures 4F and 4G**). To confirm that slower/less efficient SV retrieval was due to mitochondrial dysfunction, and not caused by other synaptic defects, we supplied Sirt3 KD terminals with glucose instead of lactate and pyruvate. Simply bypassing OXPHOS with glycolysis was sufficient to restore SV retrieval in Sirt3 KD terminals, validating the functional integrity of synapses that lack Sirt3 (**Figures 4F and G**).

Sirt3 has been implicated in buffering cytosolic Ca^2+^ during excitotoxicity^18^. Since Ca^2+^ modulates SV endocytosis^41^, we sought to confirm that the SV retrieval defect in Sirt3 KD neurons is not secondary to impaired Ca^2+^ dynamics. To this end, we examined cytosolic Ca^2+^dynamics during electrical activity using physin-GCaMP6f in which the Ca^2+^ sensor GCaMP6f is anchored at nerve terminals. Sirt3 KD had no effect on presynaptic Ca^2+^ spikes evoked by a single AP or a short train of 10 APs (**Figures S2E-S2H**), suggesting that Sirt3 depletion does not alter cytosolic Ca^2+^ buffering under physiological levels of synaptic activity. Altogether, we conclude that Sirt3 sustains SV retrieval in glucose-deprived nerve terminals by stimulating mitochondrial ATP synthesis.

## Discussion

Glucose is the preferred fuel for the brain^42^, yet its availability varies in time and space within various brain regions, particularly during intense circuit activity, or dietary restrictions^5^. Thus, compensatory metabolic pathways are essential for sustaining neuronal function when glucose availability becomes limited. Here, we show that neuronal glucose deprivation activates the CREB transcriptional program and induces the expression of the mitochondrial master regulator PGC1α, and its target gene, the mitochondrial deacetylase, Sirt3. Furthermore, we demonstrate that Sirt3 stimulates mitochondrial oxidative capacity for ATP production in nerve terminals and provides energetic support for synaptic transmission in the absence of glucose. Our findings are consistent with the emerging evidence for the metabolic flexibility of neurons in utilizing alternative energy sources^43^, particularly lactate, a circulating metabolite that fuels many organs^44^. While mitochondrial Ca^2+^ plays a critical role in rapid stimulation of oxidative metabolism during electrical activity^20^, we found that the prolonged metabolic shift to lactate/pyruvate consumption is mediated by transcriptional and post-translational modulation of mitochondrial function. The activation of CREB signaling during glucose deprivation is noteworthy given its well-established role in learning and memory^45, 46^, which are also energetically intensive processes^47, 48^. It is tempting to speculate that the activation of CREB signaling during memory formation results from local hypoglycemia caused by sustained neuronal activity.

PGC1α is a transcriptional coactivator with many downstream targets, including Sirt3. Here we demonstrated a role for basal Sirt3 in regulating mitochondrial oxidation of pyruvate in nerve terminals. Furthermore, we show that the induction of Sirt3 expression by the CREB-PGC1α signaling directly links the transcriptional response to glucose deprivation with post-translational modulation of mitochondrial function via lysine deacetylation (**Figure 2**). At present, it is unclear whether Sirt3 activation occurs in a cell-wide or compartment-specific manner. However, localized modulation of Sirt3 activity can be mediated by changes in the mitochondrial NAD^+^ content, as Sirt3 is an NAD^+^-dependent deacetylase, and neuronal NAD^+^/NADH ratio fluctuates significantly during electrical activity^49^. Therefore, future studies are needed to investigate if and how mitochondrial NAD^+^ levels in nerve terminals regulate Sirt3 activity, particularly during glucose deprivation or electrical activity.

In nerve terminals, the SV cycle is one of the most energy consuming processes of synaptic transmission^38, 50^. The SV cycle includes multiple events, such as the release of SV though fusion with the plasma membrane, SV retrieval through endocytosis, and re-acidification and refilling of SVs with neurotransmitter. While the retrieval step of the SV cycle has been known to be most susceptible to energetic imbalances caused by mitochondrial or glycolytic inhibition^20, 38, 51^, specific metabolic regulators, such as Sirt3, were not previously identified (**Figures 4F and 4G**). The greater susceptibility of SV retrieval to ATP levels is often attributed to the energy requirements of dynamin (in the form of GTP) to pinch off vesicles during endocytosis^52–54^. In contrast, the release of SVs via exocytosis is not impacted by low ATP in Sirt3-deficient nerve terminals (**Figure 4E**), suggesting that the docking and release of SVs have a lower energetic barrier than SV retrieval. Our findings reinforce the idea that different cellular processes have distinct ATP thresholds and are differentially susceptible to metabolic perturbations in disease states.

Here, we elucidated the function of Sirt3 in hippocampal (excitatory) synapses, yet Sirt3 may regulate synaptic transmission differently in other types of neurons, such as inhibitory interneurons, which have distinct activity patterns and metabolic demands. Although our primary hippocampal cultures contain a mixture of neurons and glia (mainly astrocytes), we do not know what role, if any, Sirt3 plays in astrocytic metabolism or the metabolic cross-talk between neurons and astrocytes. Given the critical function of glia in regulating synaptic function and plasticity^55^, future studies are needed to uncover the glial functions of Sirt3. Our characterization of Sirt3 in mammalian synapses is relevant to many disease states, such as obesity and aging, where Sirt3 expression declines significantly^56, 57^. Indeed, our findings suggest that neuronal Sirt3 deficiency may contribute to synaptic dysfunction or cognitive impairments in these conditions. Therefore, it will be important to determine whether restoring Sirt3 activity in the brain is therapeutic in these and other disease states.

---

**Figure S1. (Related to Fig 2.).**
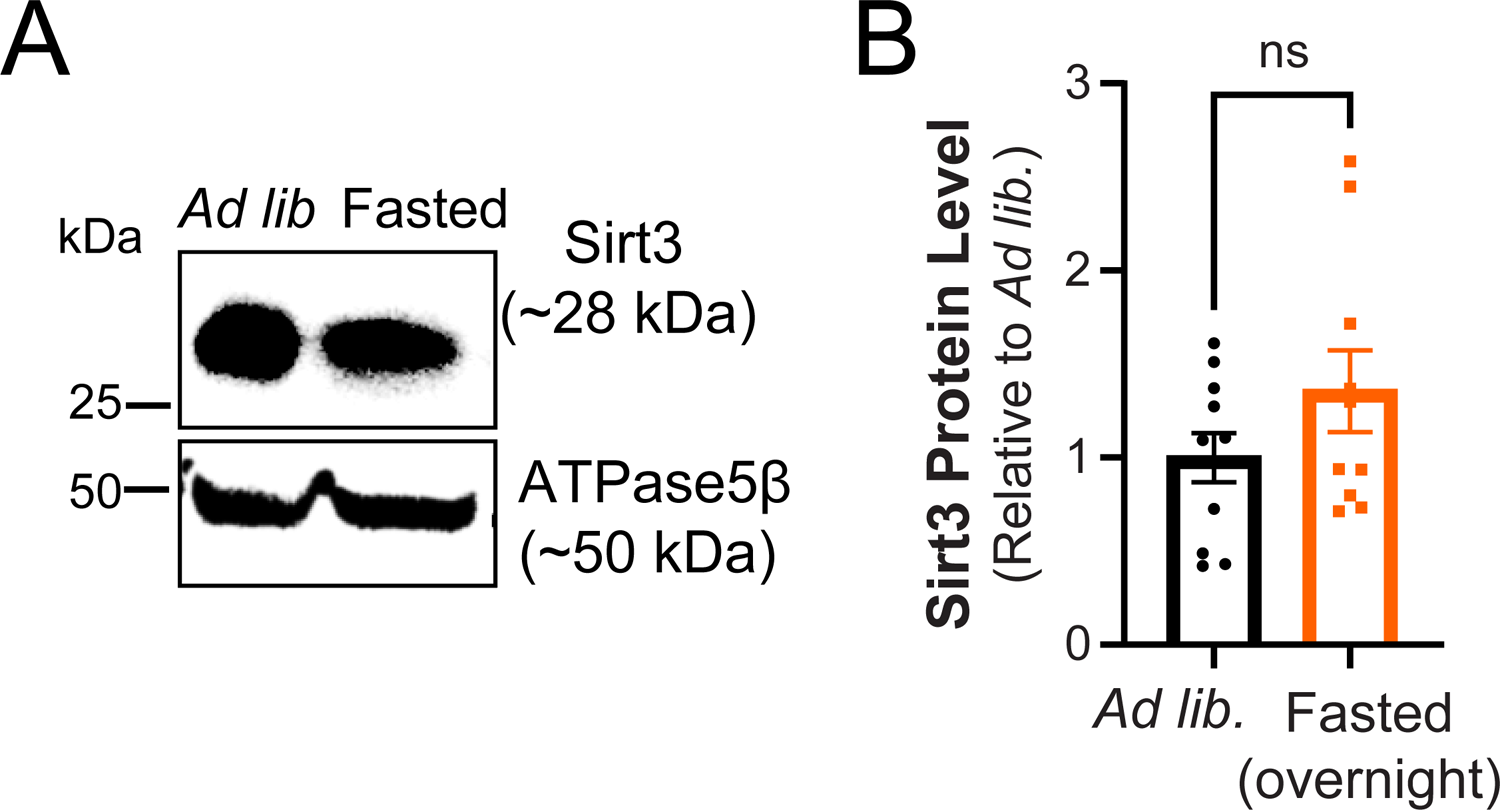
Hippocampal Sirt3 expression is not altered by overnight fasting. **A)** Hippocampal tissues from mice fed *ad lib*. or fasted overnight were immunoblotted for Sirt3 and mitochondria ATPase5β. **B)** Sirt3 band intensity normalized to ATPase5β and expressed relative to *ad lib* samples. n = 11 animals. Average normalized Sirt3 band intensity ± SEM: *ad lib.*, 1.00 ± 0.13; overnight fasted, 1.67 ± 0.36. n.s., not statistically significant. Unpaired t-test.

**Fig. S2. (Related to Fig. 4).**
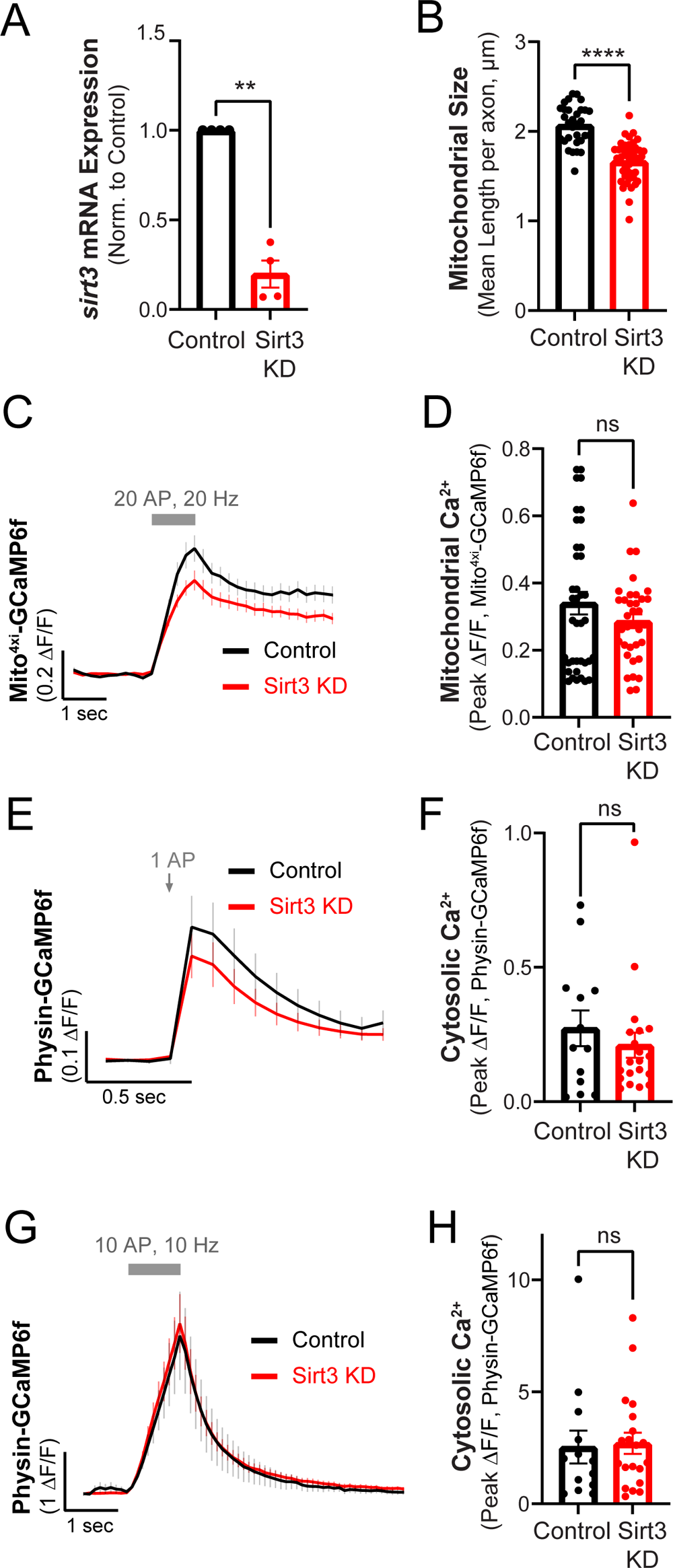
The impact of Sirt3 KD on cytosolic and mitochondrial calcium dynamics, and mitochondrial size in axons. A) Relative expression of *sirt3* mRNA in control and Sir3 KD cortical neurons. Values are normalized to *β-actin* mRNA and expressed relative to the control. n= 4 cortical samples. Average normalized mRNA level ± SEM: Sirt3 KD, 0.19 ± 0.07. B) Mitochondria length in control and Sirt3 KD axons. Mean length per axon (μm) ± SEM: control, 2.0 ± 0.03, Sirt3 KD, 1.6 ± 0.03 n= 31-42 neurons. C) Average traces of mito^4xi^-GCaMP6f showing mitochondrial Ca^2+^ uptake in control and Sirt3 KD axons stimulated with 20 AP at 20 Hz. D) Peak responses of Mito^4xi^ - GCaMP6f (ΔF/F) following stimulation. n = 26-38 neurons. mean ΔF/F ± SEM: control, 0.34 ± 0.03, Sirt3 KD, 0.28 ± 0.02. E) Average traces of physin-GCaMP6f showing cytosolic Ca^2+^ uptake in control and Sirt3 KD axons stimulated with 1AP. F) Peak responses of physin-GCaMP6f (ΔF/F) to stimulation. n = 13-20 neurons, mean ΔF/F ± SEM: control, 0.27 ± 0.06, Sirt3 KD, 0.21 ± 0.04. G) Average traces of physin-GCaMP6f showing cytosolic Ca^2+^ uptake in control and Sirt3 KD axons stimulated with 10 AP at 10 Hz. H) Peak responses of physin-GCaMP6f (ΔF/F). n = 13-20 neurons, mean ΔF/F ± SEM: control, 2.5 ± 0.7, Sirt3 KD, 2.7 ± 0.4. Unpaired t-test (A, B, D, F, H). Error bars are SEM.

## Acknowledgements

We thank the Animal Surgery Core at Hope Center for Neurological Disorders, and the Washington University Center for Cellular Imaging, and the Genome Technology Access Center at the McDonnell Genome Institute at Washington University School of Medicine Center, which is partially supported by NCI Cancer Center Support Grant #P30 CA91842 to the Siteman Cancer Center and by ICTS/CTSA Grant# UL1TR002345 from the National Center for Research Resources (NCRR), a component of the National Institutes of Health (NIH), and NIH Roadmap for Medical Research. We thank T. Roostaei and A. Azeri for their help with RNA sequencing analysis, and B. Finck for anti-MPC1 antibody and PGC1α adenovirus. This work was funded by the Washington University Institute of Clinical and Translational Sciences which is, in part, supported by the NIH/National Center for Advancing Translational Sciences (NCATS), CTSA grant #UL1 TR002345, the McDonnell Institute for Cellular Neurobiology Small Grant Program (G.A.), the Klingenstein-Simons Foundation (G.A.), the Whitehall Foundation (G.A.), NIGMS R35GM147222 (G.A.), and NINDS R35 NS111596 (V.K.).

## Author Contributions

Conceptualization, G.A.; Investigation, A.T., A.H., M.L., D.M, T.H., M.S., and G.A.; Writing – Original Draft, A.T., G.A.; Writing – Review & Editing, A.T., V.K., G.A., Visualization, A.T, D.M., and G.A.; Funding Acquisition, V.K., C.B., and G.A; Supervision, G.A.

## STAR METHODS

### Resource Availability

#### Lead Contact

Further information and requests for resources and reagents should be directed to and will be fulfilled by the Lead Contact Ghazaleh Ashrafi at.

#### Material Availability

Plasmids generated in this study have been deposited to addgene.

#### Data and Code Availability

RNA-seq data will be deposited to the GEO repository. Syn-ATP analysis code is available on GitHub.

### Experimental Model and Subject Details

#### Animals

Animal-related experiments were performed with wild-type rats of the Sprague-Dawley strain (Charles River code 400, RRID: RGD_734476) in accordance with protocols approved by the Washington university IACUC.

In intermittent fasting experiments, hippocampi were dissected from female C57BL/6 mice fed *ad libitum* or intermittent fasted on 24-hour cycles for 6 months, starting on 6 weeks of age. All animals were placed on aspen bedding for the duration of study and cages were changed for both mouse groups on fasting days mice fed ad libitum. Intermittently fasted mice were sacrificed during the fasting phase. In overnight fasting experiments, hippocampi were dissected from 11-week old C57BL/6 male mice that were fed *ad libitum* or fasted overnight for 12 hours.

#### Hippocampal neuronal cultures

Hippocampi were dissected from 1-3 days old rat pups of mixed sex from the Sprague-Dawley strain. Tissues were dissociated and mixed cultures of neurons and glia were plated on coverslips coated with poly-ornithine and transfected 6-8 days after plating with calcium phosphate as previously described^1^. Hippocampal neurons were maintained in culture media composed of MEM (Thermo Fisher Scientific 51200038), 0.6% glucose, 0.1 g/L bovine transferrin (Millipore 616420), 0.25 g/L insulin, 0.3 g/L GlutaMAX^TM^ supplement (Thermofisher 35050-061), 5% fetal bovine serum (R&D Systems S11510), 2% N-21 (R&D Systems AR008), and 4 μm cytosine β-d-arabinofuranoside (Ara-C), added after 2-3 days *in vitro* (DIV) to limit glial proliferation. Cortical neurons were maintained in Neurobasal-A media (NB-A) (Gibco # A24775-01) supplemented with 2% N-21, 5% fetal bovine serum, and 4 μm Ara-C (added after 1 DIV). Cultures were incubated at 37°C in a 95% air and 5% CO_2_ humidified incubator for 10 - 14 DIV (cortical neurons) or 14 - 21 DIV (hippocampal neurons) prior to use.

## Method Details

### Plasmid Constructs

The following previously published DNA constructs were used: vGLUT1-pHluorin^2^, Syn-ATP^3^,, (synapto-)physin-GCaMP6f^4^, mito^4x^-RFP (gift of T.A. Ryan). The mito^4xi^-GCaMP6f construct (addgene #192075) with improved mitochondrial targeting. A human Sirt3-RFP driven by the CaMKII promoter was constructed through PCR of the human Sirt3 sequence from Sirt3-FLAG (Addgene plasmid 13814)^5^ into the BamHI and AgeI sites of CaMKII-MICU3-RFP plasmid^6^ resulting in the linker sequence RPVVA joining Sirt3 and RFP sequences.

We used the pLKO.1 vector (Addgene plasmid 10878)^7^ for expression of shRNA against rat Sirt3 (target: CTTACTACATGTGGCTGAT).

### Glucose Deprivation Experiments

Cortical neurons 10-14 DIV were treated with Neurobasal-A (NB-A) media without glucose (Gibco # A24775-01) or with media supplemented with 5-mM glucose (control condition). After 1-3 hours of incubation at 37°C, the cells were lysed for either RNA or protein extraction or immunostaining.

### RNA Isolation and Quantitative PCR

Total RNA was isolated from primary dissociated cortical neuron cultures using RNeasy Mini Kits (QIAGEN 74104). cDNA was synthesized by reverse transcription using All-In-One 5X RT Master Mix (ABM-G592) following manufacturer’s protocol. qPCR was set up with cDNA as template and PowerUp™ SYBR™ Green Master Mix (Thermo Fisher, A25742) or TaqMan® Fast Advanced Master Mix (Applied Biosytems, 4444557). Relative mRNA expression was quantified by normalizing to housekeeping genes (*Gapdh*, *18s* rRNA or *β-actin*) using ΔΔC_t_ method.

### Production and Application of Lentiviral Particles

Lentivirus were produced by the Hope Center Viral Vectors Core at Washington University School of Medicine. Viral aliquots were functionally titrated and the exact volume of lentivirus required to achieve maximal transduction of hippocampal or cortical cultures was determined using BFP expression, driven by a CMV promoter on the pLKO.1 backbone. For imaging experiments, lentivirus was added to neurons 5 DIV and after 2 days of infection, media exchange was performed. All experiments were performed at least 7 days after viral transduction to maximize knockdown efficiency.

### Western Blotting

Cultured cortical cells and brain tissues were solubilized in the neuronal extraction buffer (, Thermo Fisher Scientific, 87792) supplemented with 1X protease inhibitor cocktail and 0.1 mM PMSF. Protein concentrations were in quantified using s BCA assay kit (BioVision) and a BioTek Synergy plate reader and Gen5 software (BioTek Instruments, Winooski, VT). Immunoblotting was conducted using 4–10% SDS gradient polyacrylamide gel followed by transfer on PVDF membrane. The following primary antibodies were used Sirt3 (1:1000, Cell Signaling Technology 5490S) anti-acetyl lysine (1:1000, Cell Signaling Technology 9441S), ATPase5β (1:1000, Sigma-Aldrich HPA001520), β-actin (1:5000, BioRad HCA147P), α-tubulin (1:2000, Sigma-Aldrich T6074).

### Mitochondria Fractionation

Cultured cortical cells were homogenized in mitochondria isolation buffer (10mM Tris-HCl, pH 7.4, 1mM EDTA, pH 8.0, 250 mM Sucrose, 10 mM KCl, 1x Protease inhibitor, 0.1 mM PMSF and deacetylase inhibitor (1;200, Santa Cruz sc-362323) and centrifuged at 4°C for 10 minutes. The supernatant was collected and centrifuged for 20 minutes at 4°C. The mitochondrial pellet was washed thrice with mitochondrial isolation buffer and the supernatant containing the cytosolic fraction was concentrated using Amicon Ultra 0.5 centrifugal filter devices. The mitochondrial and cytosolic samples were prepared using 1x Lamelli buffer (BioRad, 1610747).

### Immunofluorescence and Confocal Microscopy

Hippocampal neurons were fixed with 4% PFA, permeabilized with 0.5 % Triton, blocked with 5% bovine serum albumin (BSA) for 1 hour at room temperature (RT), and incubated at room temperature for 2 hours or overnight at 4°C with the following primary antibodies: Sirt3 (1:200, Cell Signaling Technology 5490S), TOMM20 (1:500, Cell Signaling Technology 42406S), ATPase5β (1:500, Sigma HPA001520), Tuj1 (1:1000, R&D Systems MAB1195), pCREB (Cell Signaling Technology, 9198S). Coverslips were then incubated with the following secondary antibodies: anti-rabbit Alexa-fluor568 (1:500, Thermo Fisher A21428) and anti-mouse Alexa-fluor488 (1:500, Thermo Fisher A11059) for 1 hour at RT. For staining of nuclei Hoechst (1:2000, Invitrogen H3570) was used at RT for 10 min and coverslips were then mounted with anti-fade mounting media (Thermo Fisher P36965), and kept at 4°C until imaged. Confocal images were collected on a Zeiss LSM 880 Confocal Microscope at the Washington University Center for Cellular Imaging which was purchased with support from the Office of Research Infrastructure Programs (ORIP), as part of the NIH Office of the Director under grant OD021629.

### Live Imaging of Neurons

Imaging experiments were performed on a custom-built laser illuminated epifluorescence microscope with an Andor iXon Ultra 897 camera. Coverslips were mounted in a laminar flow perfusion chamber and perfused with Tyrodes buffer (pH 7.4) containing (in mM) 119 NaCl, 2.5 KCl, 2 CaCl_2_, 2 MgCl_2_, 50 HEPES, 5 glucose or 1.25 lactate and 1.25 pyruvate, supplemented with 10 μM 6-cyano-7nitroquinoxalibe-2, 3-dione (CNQX), and 50 μM D,L-2-amino-5phosphonovaleric acid (APV) (both from Sigma-Aldrich) to inhibit post-synaptic responses. Action potentials were evoked in neurons with 1 ms pulses creating field potentials of ∼10 V/cm via platinum–iridium electrodes. Temperature was maintained at 37°C using an Okolab stage top incubator.

Synaptic vesicle endocytosis in nerve terminals of hippocampal neurons was studied using vGLUT1-pHluorin as previously described by Ashrafi et al., 2020. Neurons were stimulated with trains of 100 AP at 10Hz and images were acquired at 2Hz. For alkalization of pHluorin-containing vesicles, NH_4_Cl solution was used which had a similar composition as Tyrodes buffer except it contained 50 mM NH_4_Cl and 66.5 mM NaCl. Cytosolic Ca^2+^ signaling experiment was done using synGCaMP6f in which GCaMP6f is coupled with synaptophysin as described earlier (de Jaun-Sanz et al., 2017, Brockhaus et al., 2018). Neurons were stimulated with trains of 10 AP first at 10Hz and then 1Hz and data was acquired at 10Hz imaging. Mitochondrial Ca^2+^ signaling experiments using mito^4x^-GCaMP6f were performed as described^6^. Neurons were stimulated with trains of 20 AP at 20Hz and images were acquired at 5Hz. As a standard, 30 image frames were recorded before the stimulus train was triggered.

### Quantification and Statistical Analysis

#### Image Analysis and Statistics

Images were analyzed using the ImageJ plugin Time Series Analyzer where ∼20-50 regions of interest (ROIs) of ∼2 μm or corresponding to responding synaptic boutons were selected and the fluorescence was measured over time. All fitting was done with GraphPad Prism v9.0 for Windows. We used nonparametric Mann–Whitney U test to determine the significance of the difference between two unpaired conditions, p < 0.05 was considered significant and denoted with a single asterisk, whereas p < 0.01, p < 0.001 and p < 0.0001 are denoted with two, three, and four asterisks, respectively.

#### Quantification of nuclear phospho-CREB intensity

Analysis of phosph-CREB immunostaining of neuronal cultures was performed with ImageJ. Briefly, ROIs corresponding to Hoechst-stained nuclei (7-500 μm in size) were selected using particle analysis. The ROIs were transferred to the FITC channel corresponding to pCREb staining and average fluorescence intensity was calculated. The pCREB intensity was then normalized to the mean of control and plotted using GraphPad prism v9.0.

#### Quantification of presynaptic ATP level

Luminescence imaging of the presynaptic ATP reporter, Syn-ATP, and image analysis was performed using a semi-automatic platform, as previously described^8^. All measurements were pH corrected as previously described^3^.

#### Quantitative analysis of release probability

vGLUT1-pH responses in single hippocampal terminals were recorded by stimulating cultures at 1Hz for 100 seconds, followed by 100AP at 10Hz, separated by a 10-second rest period. Single fusion events were determined during the 1Hz period using a Matlab custom code^9^ based on the change in fluorescence. Release at high frequency was calculated, like the relation between total fluorescence change during the 100Hz stimulation period and the quantum size of a single synaptic vesicle fusion event. The quantal size was determined as the average fluorescence change of the events detected at the same synapsis at 1Hz, a minimum of five single events was required for reliable quantification.

#### Quantification of synaptic vesicle retrieval block

Endocytic time constants (τ) were calculated by fitting the fluorescent change after the stimulus to a single exponential decay as described earlier^10^. The fractional retrieval block in endocytosis of vGLUT1-pH was calculated as the fraction of ΔF remaining after 2 times the average endocytic time constant of the control (2τ), divided by the maximum ΔF at the end of stimulation (ΔF_2τ_/ ΔF_max_).

#### Mitochondrial and cytosolic Ca^2+^ measurements

Mito^4xi^-GCaMP6f was used for measuring mitochondrial Ca^2+^ dynamics and data were obtained from imaging at 5 Hz from an average of 3 trials. For cytosolic Ca^2+^ measurements at 1 Hz with 10AP, data was acquired at 10Hz imaging and the average peak was calculated by averaging the ΔF values after every 0.1s for the entire duration of stimulus (2.8 to 3.8s). physin-GCaMP and Mito^4xi^-GCaMP6f peak fluorescence (Δ*F/F*_0_) was found by averaging the 3 highest values of Δ*F/F*_0._

#### Quantification of mitochondrial length in neuronal axons

Hippocampal neurons expressing mito**^4x^**-RFP with or without Sirt3 shRNA were fixed and imaged with confocal microscopy. ImageJ was used to threshold images and measure mitochondrial size in neuronal axons.

#### RNA Sequencing and Analysis

Cortical neuronal cultures 13 DIV were treated with NB-A media containing 5mM glucose (control) or NB-A without glucose (glucose deprivation) for 3 hours. Total RNA was isolated using RNeasy Mini Kits (QIAGEN 74104) and triplicate samples were submitted to the GTAC core at McDonnell Genome Institute, Washington University. Samples were prepared according to library kit manufacturer’s protocol, indexed, pooled, and sequenced on an Illumina NovaSeq 6000. Basecalls and demultiplexing were performed with Illumina’s bcl2fastq software and a custom python demultiplexing program with a maximum of one mismatch in the indexing read. RNA-seq reads were then aligned to the Ensembl release 76 primary assembly with STAR version 2.5.1a^11^. Gene counts were derived from the number of uniquely aligned unambiguous reads by Subread:featureCount version 1.4.6-p5^12^. Isoform expression of known Ensembl transcripts were estimated with Salmon version 0.8.2^13^. Sequencing performance was assessed for the total number of aligned reads, total number of uniquely aligned reads, and features detected. The ribosomal fraction, known junction saturation, and read distribution over known gene models were quantified with RSeQC version 2.6.2^14^.

All gene counts were then imported into the R/Bioconductor package EdgeR^15^ and TMM normalization size factors were calculated to adjust for samples for differences in library size. Ribosomal genes and genes not expressed in the smallest group size minus one samples greater than one count-per-million were excluded from further analysis. The TMM size factors and the matrix of counts were then imported into the R/Bioconductor package Limma^16^. Weighted likelihoods based on the observed mean-variance relationship of every gene and sample were then calculated for all samples and the count matrix was transformed to moderated log 2 counts-per-million with Limma’s voomWithQualityWeights^17^. The performance of all genes was assessed with plots of the residual standard deviation of every gene to their average log-count with a robustly fitted trend line of the residuals. Differential expression analysis was then performed to analyze for differences between conditions and the results were filtered for only those genes with Benjamini-Hochberg false-discovery rate adjusted p-values less than or equal to 0.05.

For each contrast extracted with Limma, global perturbations in known Gene Ontology (GO) terms, MSigDb, and KEGG pathways were detected using the R/Bioconductor package GAGE^18^ to test for changes in expression of the reported log 2 fold-changes reported by Limma in each term versus the background log 2 fold-changes of all genes found outside the respective term. The R/Bioconductor package heatmap3^19^ was used to display heatmaps across groups of samples for each GO or MSigDb term with a Benjamini-Hochberg false-discovery rate adjusted p-value less than or equal to 0.05. Perturbed KEGG pathways where the observed log 2 fold-changes of genes within the term were significantly perturbed in a single-direction versus background or in any direction compared to other genes within a given term with p-values less than or equal to 0.05 were rendered as annotated KEGG graphs with the R/Bioconductor package Pathview^20^.

To find the most critical genes, the Limma voomWithQualityWeights transformed log 2 counts-per-million expression data was then analyzed via weighted gene correlation network analysis with the R/Bioconductor package WGCNA^21^. Briefly, all genes were correlated across each other by Pearson correlations and clustered by expression similarity into unsigned modules using a power threshold empirically determined from the data. An eigengene was then created for each de novo cluster and its expression profile was then correlated across all coefficients of the model matrix. Because these clusters of genes were created by expression profile rather than known functional similarity, the clustered modules were given the names of random colors where grey is the only module that has any pre-existing definition of containing genes that do not cluster well with others. These de-novo clustered genes were then tested for functional enrichment of known GO terms with hypergeometric tests available in the R/Bioconductor package clusterProfiler^22^. Significant terms with Benjamini-Hochberg adjusted p-values less than 0.05 were then collapsed by similarity into clusterProfiler category network plots to display the most significant terms for each module of hub genes in order to interpolate the function of each significant module. The information for all clustered genes for each module were then combined with their respective statistical significance results from Limma to determine whether or not those features were also found to be significantly differentially expressed.

Analysis of transcription factor target genes in Figure 1 was performed on differentially expressed genes with p-values <0.01 using the FUMA GWAS (Functional Mapping and Annotation of Genome-Wide Association Studies) platform^23^.

## KEY RESOURCES TABLE

| REAGENT or RESOURCE | SOURCE | IDENTIFIER |
| --- | --- | --- |
| **Antibodies** |  |  |
| Guinea pig anti-vGLUT1 | Millipore Sigma | CAT#: AB5905, |
| anti-Tuj1 (β-tubulin III) antibody | R&D Systems | MAB1195 |
| anti-ATPase5β antibody | Sigma-Aldrich | CAT#: HPA001520 |
| anti-Sirt3 antibody | Cell Signaling Technology | CAT#: 5490S |
| anti-β actin antibody | Bio-Rad | CAT#: HCA147P |
| anti-acetylated Lysine antibody | Cell Signaling Technology | CAT#: 9441S |
| anti-α tubulin antibody | Sigma-Aldrich | CAT#: T6074 |
| anti-TOMM20 antibody | Cell Signaling Technology | CAT#: 42406S |
| anti-Flag antibody | Sigma- Aldrich | CAT#: F1804 |
| anti-pCREB antibody | Cell Signaling Technology | CAT#: 9198S |
| **Biological Samples** |  |  |
| rSirt3 shRNA Lentivirus | Hope Center Viral Vectors<br>Core at Washington<br>University in St. Louis | N/A |
| PGC1α AAV | Gift of B. Finck | N/A |
| GFP AAV | Gift of B. Finck | N/A |
| **Chemicals, Peptides, and Recombinant Proteins** |  |  |
| Dorsomorphin | Sigma-Aldrich | CAT#: 866405-64-3 |
| Deacetylation Inhibition Cocktail | Santa Cruz | CAT#: sc-362323 |
| 6-cyano-7nitroquinoxaline-2, 3-dione (CNQX) | Sigma-Aldrich | CAT#: C239-25MG |
| D,L-2-amino-5phosphonovaleric acid (APV) | Sigma-Aldrich | CAT#: A5282 |
| **Critical Commercial Assays** |  |  |
| All-In-One 5X RT Master Mix | Applied Biosystems | CAT#: ABM-G592 |
| PowerUp™ SYBR™ Green Master Mix | Thermo Fisher | CAT#: A25742 |
| TaqMan® Fast Advanced Master Mix | Applied Biosystems, | CAT#: 4444557 |
| **Deposited Data** |  |  |
| RNA-seq data | The GEO repository |  |
| Syn-ATP analysis code | GitHub |  |
| **Experimental Models: Cell Lines** |  |  |
| **Experimental Models: Organisms/Strains** |  |  |
| Sprague-Dawley Rat | Charles River | Strain code: CD IGS<br>SD # 001<br>RRID: RGD_734476 |
| C57BL/6NJ Mice (overnight fast) | Jackson Labs | Strain code: 005304<br>RRID:IMSR_JAX:00<br>5304 |
| C57BL/6J Mice (alternate-day fast) | Jackson Labs | Strain code: 000664<br>RRID:IMSR_JAX:00<br>0664 |
| **Recombinant DNA** |  |  |
| vGLUT1-pHluorin | Voglmaier et al., 2006 | N/A |
| Syn-ATP | Rangaraju et al., 2014 | Addgene #51819 |
| synaptophysin-GCaMP6f | de Juan-Sanz et al., 2017 | N/A |
| mito<sup>4x</sup>-RFP | gift of T.A. Ryan | Addgene #192075 |
| mito<sup>4xi</sup>-GCaMP6f | Ashrafi et al., 2020 | Addgene #192075 |
| pLKO.1 vector | Moffat et al., 2006 | Addgene #10878 |
| hSirt3-RFP | This paper |  |
| Sirt3-FLAG | North et al., 2003 | Addgene #13814 |
| **Sequence-Based Reagents** |  |  |
| shRNA targeting rSirt3 | This paper |  |
| **Software and Algorithms** |  |  |
| ImageJ | National Institute of Health | https://imagej.nih.gov/ij/ |
| GraphPad Prism 9.2.0 | GraphPad Software | http://www.graphpad.com/scientific-software/prism/ |
| OriginPro 9 | OriginLab | http://www.originlab.com/ |
| Jupyter Notebook | Project Jupyter | https://jupyter.org/ |
| Adobe Illustrator | Adobe Inc. | www.adobe.com |
| **Other** |  |  |

